# Tumour marker analysis using a machine learning assisted vibrational spectroscopy approach

**DOI:** 10.64898/2026.03.27.714840

**Authors:** Rashad Fatayer, Stephen John Sammut, Ganapathy Senthil Murugan

**Affiliations:** Optoelectronics Research Centre, University of Southampton, Southampton, UK; The Breast Cancer Now Toby Robins Research Centre, The Institute of Cancer Research, London, UK; The Royal Marsden Hospital NHS Foundation Trust, London, United Kingdom

## Abstract

Tumour biomarkers such as CA125, CA15-3, CA19-9, AFP and CEA are routinely used in the oncology clinic to diagnose cancer, monitor response to therapy, and detect relapse. However, their quantification depends on immunoassay-based methods that are time-consuming, reagent-dependent, and poorly suited to resource-limited settings. Here, we present a machine learning-assisted ATR-FTIR spectroscopy approach for label-free tumour biomarker analysis to enable simple and rapid quantification at the bedside. Using principal component analysis (PCA), we first demonstrate that these five clinically relevant biomarkers are spectrally separable, with the protein-associated region (1200-1700 cm^−1^) providing the greatest discriminative information. We then develop partial least squares regression (PLSR) models to quantify CA125 in phosphate-buffered saline (R^2^ = 0.95) and in human serum across a clinically relevant concentration range, achieving reliable predictions at and above the clinical decision threshold of 35 U/mL. A semi-quantitative classification model further demonstrated robust identification of elevated CA125, with a macro-average sensitivity of 0.86 and specificity of 0.92. These results support ATR-FTIR spectroscopy as a rapid, reagent-free platform for cancer biomarker monitoring, with potential utility in resource-limited settings.

## Introduction

Cancer remains a leading global health challenge, accounting for approximately one in six deaths worldwide, with an incidence projected to rise to 35 million new cases by 2050 ^1^. This growing burden places substantial pressure on healthcare systems, particularly in resource-constrained settings. Because patient outcomes depend strongly on timely and accurate clinical decision-making across the cancer care pathway ^2^, there is growing interest in the use of tumour biomarkers that can inform early detection, treatment response monitoring, disease surveillance, and prognostic assessment ^3^.

Protein based tumour biomarkers represent a major class of biomarkers used in the clinic. These proteins enter the circulation through tumour cell secretion ^4^, packaging into extracellular vesicles (EVs) ^5^, or following cell death ^6^. Widely utilised examples of these protein biomarkers include carbohydrate antigen 125 (CA125) in ovarian cancer ^7^, carbohydrate antigen 15-3 (CA15-3) in breast cancer ^8^, carbohydrate antigen 19-9 (CA19-9) in pancreatic cancer ^9^, alpha-fetoprotein (AFP) in hepatocellular carcinoma and testicular cancer ^10^, and carcinoembryonic antigen (CEA) in colorectal cancer ^11,12^. These biomarkers are routinely used in oncological practice to aid diagnosis and surveillance of cancer, with clinical interpretation based on measured blood concentrations relative to established thresholds ^13^.

Current quantification methods rely on immunoassay-based methods (e.g. ELISA derived, chemiluminescent, electrochemiluminescent, and fluorescent immunoassays ^14^), which involve multiple processing steps, including incubation, washing, and separation, and are therefore resource intensive^15^. Moreover, the dependence on specialised reagents and controlled storage, alongside the need for dedicated laboratory infrastructure, increases cost and restricts deployment outside central laboratories ^16^. As a result, access can be limited in resource-constrained settings. Taken together, these limitations highlight the need for faster and low-cost analytical approaches that are suitable for decentralised or point-of-care usage.

In recent years, vibrational spectroscopy, particularly attenuated total reflectance-Fourier transform infrared (ATR-FTIR) spectroscopy, has offered an alternative that addresses these requirements. The technique enables rapid, label-free analysis of biological samples using portable instrumentation with minimal sample preparation. In ATR-FTIR, infrared radiation is absorbed by molecular bonds within the sample, producing characteristic spectral fingerprints that reflect biochemical composition ^17^. This capability has shown potential across various cancer diagnostic applications and tumour types ^18–21^. However, most studies to date have predominantly focused on qualitative detection tasks, such as binary classification of disease presence or absence, with comparatively limited attention given to quantitative biomarker measurement. Consequently, a translational gap remains between proof-of-concept ATR-FTIR demonstrations and clinical workflows, where quantitative outputs are needed to apply diagnostic thresholds, support longitudinal monitoring, and assess treatment response.

To address this gap, we have developed a machine learning assisted ATR-FTIR spectroscopy methodology for protein tumour biomarker analysis. First, we assessed the spectral separability of five biomarkers currently used in the clinic (CA125, CA15-3, CA19-9, AFP, and CEA) using unsupervised multivariate analysis of spectra acquired from samples prepared in phosphate-buffered saline (PBS), demonstrating that these biomarkers produce distinct spectral features within protein-associated regions. Building on this foundation, we developed and validated a quantitative regression model for CA125 at high concentrations in phosphate-buffered saline (PBS), before extending the approach to physiologically relevant concentrations and human serum samples. To our knowledge this study is the first to systematically show that machine learning assisted ATR-FTIR spectroscopy can achieve quantitative cancer biomarker measurements aligned with clinical requirements, offering a potential pathway toward rapid, reagent-free, and instrument-portable cancer biomarker monitoring, particularly suited to laboratory resource-limited settings.

## Materials and Methods

### Cancer biomarker samples

To assess biomarker discrimination, a panel of cancer-associated proteins was selected. CA125 (catalogue no. P251-4), CA15-3 (catalogue no. P301-4), and CA19-9 (catalogue no. P291-4) were obtained from BBI Solutions as frozen solutions derived from human carcinoma cell lines and supplied as single, homogeneous batches in phosphate buffered saline (PBS). The supplied concentrations were 322 kU/mL for CA125, 19.8 kU/mL for CA15-3, and 204 kU/mL for CA19-9, as confirmed by the Roche Modular immunoassay. PBS buffer was additionally supplied separately by BBI Solutions for use in subsequent CA125 dilution experiments. CEA (catalogue no. HY-P70675) and AFP (catalogue no. HY-P7463) were obtained from MedChemExpress as lyophilised powders expressed in HEK293 cells.

A separate CA125 preparation (catalogue no. HY-P7696, MedChemExpress), supplied as a lyophilised powder expressed in HEK293 cells, was used solely for the serum spiking experiments. This second preparation was specifically used since its lyophilised format allowed direct reconstitution and spiking into human serum without introducing PBS buffer, thereby preserving the serum composition.

### Human serum

To evaluate biomarker detection in a biological sample, fresh human serum was obtained from a single healthy female donor (aged 63 years) through ResearchDonors (product reference 462505). A total of 24 mL was purchased and shipped via overnight courier under temperature-controlled conditions. The serum was received the following day and used immediately for testing to minimize protein degradation. Fresh rather than frozen serum was used to avoid freeze thaw cycles, which could alter protein structure and introduce spectral variability. This donor was not known to have a diagnosis of cancer, providing a negative control for our biomarker spiking study.

### Sample preparation

#### Biomarker discrimination studies

Frozen cancer biomarker solutions (CA125, CA15-3, and CA19-9) from BBI Solutions were allowed to thaw under refrigerated conditions and mixed by gentle pipetting prior to analysis at their supplied concentrations. Lyophilized CEA and AFP powders from MedChemExpress were reconstituted in accordance with manufacturer specifications. Initially, the lyophilized powders were centrifuged at 3500 rpm for 5 minutes, after which PBS buffer was added to achieve a final concentration of 100 μg/mL. The reconstituted solutions were also mixed by gentle pipetting to ensure homogeneity.

### Biomarker concentration regression in PBS

To establish the quantitative capability of ATR-FTIR spectroscopy for biomarker analysis, the CA125 stock solution (322 kU/mL) was serially diluted in PBS to generate a concentration series ranging from 5 to 50 kU/mL in 5 kU/mL increments (5, 10, 15, 20, 25, 30, 35, 40, 45, and 50 kU/mL). These concentrations were deliberately selected above physiologically relevant CA125 levels (typically <35 U/mL in healthy individuals, >35 U/mL indicating potential malignancy ^22^) to ensure strong detection and to identify concentration-dependent changes in a controlled PBS matrix. This high-concentration approach allowed the optimization of experiment parameters and machine learning model development with minimal interference from buffer components, providing a foundation before progressing to the more challenging task of quantification at clinically relevant concentrations in complex serum samples.

### Biomarker detection in spiked serum

For human serum-based analysis, the lyophilized CA125 (catalogue no. HY-P7696) from MedChemExpress was centrifuged at 3500 rpm for 5 minutes, then reconstituted directly in undiluted human serum to an initial concentration of 100 μg/mL. Concentrations were converted from μg/mL to U/mL using the specific activity of 1.74×10^5^ U/mg provided by the manufacturer. Serial dilutions were then prepared in the same serum to achieve physiologically relevant concentrations of 5, 10, 20, 35, 50, 100, 200, 500, and 1000 U/mL. The concentration range was selected to span values below and exceeding the established clinical decision threshold of 35 U/mL for CA125. Human serum was specifically obtained from a 63□year□old (postmenopausal) donor, as CA125 demonstrates higher diagnostic performance for ovarian cancer and fewer false□positive elevations from benign gynaecologic and physiological factors in postmenopausal women compared with premenopausal women ^22^.

### ATR-FTIR spectroscopy

ATR-FTIR spectra were acquired using an Agilent Cary 670 FTIR spectrometer equipped with a liquid-nitrogen-cooled MCT detector and a MIRacle single-bounce ATR accessory fitted with a ZnSe crystal. For each measurement, 3 µL of sample was pipetted onto the ATR crystal and allowed to air-dry at room temperature for approximately 15 minutes to form a uniform thin film. Spectra were collected over the 600-6000 cm^−1^ range at a spectral resolution of 4 cm^−1^, with 64 co-added scans per spectrum to improve the signal-to-noise ratio.

For each sample composition, three technical replicates were acquired, defined as three independent depositions (sample was removed and the ATR crystal was cleaned, followed by re-depositing another 3 µL aliquot and dry). Within each dried film, multiple repeat spectral acquisitions were collected to capture short-term instrumental variability. In total, nine repeat spectra were collected per sample deposition (replicate measurement) prior to proceeding to the next sample.

### Preprocessing

All spectra were pre-processed for machine-learning analysis using a consistent methodology. Initial spectral processing was performed in Peak Spectroscopy software, where raw spectra were manually baseline corrected and vector normalised over the 700-1800 cm^−1^ wavenumber range. Baseline correction was applied to compensate for drift arising from instrumental effects and scattering. Vector normalisation was subsequently used to minimise variations in spectral intensity caused by differences in sample drying, effective path length, and concentration, ensuring that further analysis is driven by relative spectral features rather than absolute intensity differences. Finally, Python was used to apply a Savitzky-Golay filter to compute second-order derivatives with smoothing, reducing spectral noise while enhancing subtle peak features.

### Serum background subtraction

For experiments involving CA125 spiked human serum, an additional preprocessing step was applied to isolate the spectral contribution of the biomarkers from the serum background. A reference spectrum of unspiked serum (0 U/mL CA125) was acquired and ratioed against each CA125 spiked serum spectrum using Peak Spectroscopy software. This removed the baseline contribution arising from serum components, leaving only the spectral signal attributable to the added CA125. Importantly, this approach also compensates for any CA125 present in the donor serum at physiological levels, ensuring that the spectral features used for modelling corresponded solely to the known spiked concentrations, rather than an unknown combination of endogenous and exogenous CA125.

### Machine learning

All analyses were performed in Python 3.11 using a Jupyter Notebook environment, with data processing and machine learning implemented using NumPy, SciPy, pandas, and scikit-learn.

### Data exploration

Principal Component Analysis (PCA) was applied to the pre-processed spectra for exploratory analysis. PCA is a dimensionality reduction technique that projects the original dataset into a latent space explained by principal components (PC) that capture the maximum variance in the data. Prior to PCA, spectra were mean-centred and scaled to unit variance to prevent dominance of high-variance wavenumber regions. PCA was performed in an unsupervised manner, with the first five principal components retained to capture the dominant sources of spectral variance. PCA was used for visualisation and comparison of spectral variation between different cancer biomarkers across selected wavenumber regions, as well as between human serum samples spiked with varying concentrations of CA125. Score plots of the first two PCs, together with 95% confidence ellipses, were used to assess clustering patterns and separation between sample groups.

### Regression modelling

Partial Least Squares Regression (PLSR) was used in this study to quantify CA125 concentrations, first in buffer and subsequently in human serum. PLSR is a multivariate statistical modelling approach that relates high-dimensional input variables to a target variable through a reduced set of latent variables (LVs) used to maximise covariance, making it well suited to spectroscopic data which is highly collinear. ATR-FTIR spectra were analysed over a defined wavenumber range following the preprocessing methodology described in Section 2.5. Model optimisation and component selection were performed using a group-wise k-fold cross-validation, with spectra grouped by replicate concentration. The number of LVs was selected based on cross-validation performance, favouring parsimonious models where performance plateaued, and final model performance was evaluated on an independent test set using the coefficient of determination (R^2^) and root mean square error (RMSE).

### Classification modelling

A multi-class classification model was developed using PCA followed by Logistic Regression (PCA-LR) to classify CA125-spiked serum samples into three concentration ranges: (5-20 U/mL), (35-100 U/mL), and (200-1000 U/mL). PCA was first applied to the preprocessed ATR-FTIR spectra, followed by multinomial logistic regression to model class assignment probabilities using the Softmax function, which transforms linear combinations of PCs into a normalised probability distribution across all classes. The dataset was partitioned using a stratified replicate aware splitting strategy, where all spectra from the same biological replicate were kept together to prevent data leakage, resulting in training (60%), validation (20%), and test (20%) sets. Hyperparameter tuning was performed for a range of PCs and the inverse regularization strengths (C parameter), jointly using a grid search on the validation set, evaluating both classification accuracy and cross-entropy loss. The model performance was evaluated on the validation set using overall accuracy, per-class sensitivity, specificity, precision, with final generalisation assessed on the independent test set.

### Limit of detection

The limit of detection (LoD) for the low-concentration CA125 model in human serum (5-50 U/mL) was estimated from the held-out test set using a calibration-curve-based approach ^23^. A linear regression of predicted CA125 concentration against true CA125 concentration was fitted to the test-set results, and the LoD was calculated as:

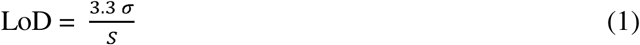

where σ is the standard deviation of the model predictions for samples at the lowest measured concentration (5 U/mL) and 5 is the slope of the regression line.

## Results and Discussion

### Overview of protein spectra

The spectral features arising from the interaction of infrared radiation with proteins followed broadly consistent trends, dominated by peptide-bond (amide) vibrations. Here, the averaged raw spectrum collected from the CA125 sample was analysed across 750-4000 cm^−1^ to identify key protein-associated features (Figure 1). The strongest contribution in the spectra was from the Amide I band occurring at ∼1660cm^−1^, mainly associated with stretching vibrations of the C=O bond, weakly coupled to C-N and N-H bonds in the peptide backbone ^24^. The next strongest feature was the Amide II band occurring at 1550cm^−1^ ^25^, attributed to N-H bending and C-N stretching. Additional protein related modes were observed in the Amide III region 1200-1400cm^−1^ which also reflected C-N stretching and N-H bending, although at lower intensities. At higher wavenumbers, the Amide A (∼3300cm^−1^) and Amide B (∼3100cm^−1^) bands were also present and are attributed to N-H stretching from hydrogen bonds. Contributions in the wavenumbers between the range of 750-1150cm^−1^ corresponded to phosphate group vibrational modes ^25^ from the PBS buffer.

**Figure 1.**
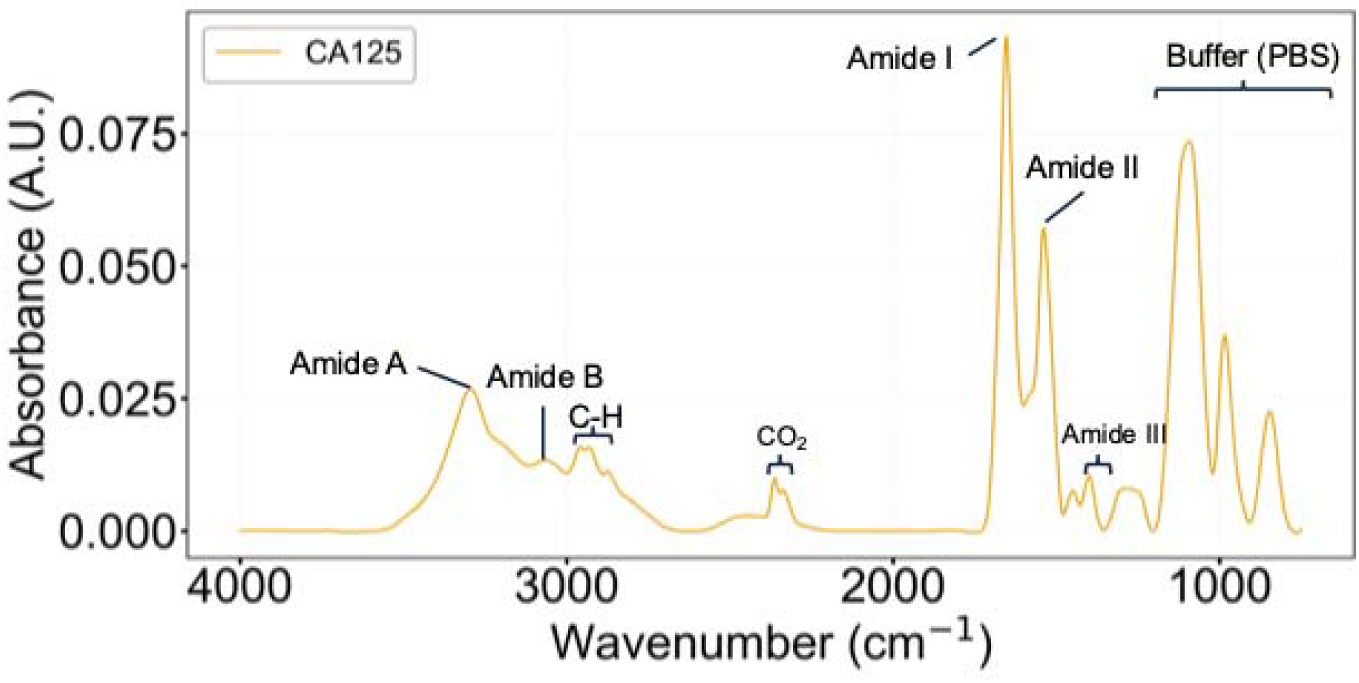
Averaged spectra of CA125 in PBS buffer.

### Comparison of major cancer biomarkers spectra

The ATR-FTIR spectra for all cancer biomarkers measured in this study, across the shared 750-4000 cm^−1^ wavenumber range, are shown in Figure 2. All biomarkers displayed protein-associated spectral features, with Amide I and Amide II bands clearly identifiable in all cases except CA19-9. CA125 and CA15-3 are peptide epitopes carried by the mucin family glycoproteins MUC16 ^26^ and MUC1 ^27^ respectively, whilst AFP and CEA belong to the oncofoetal glycoprotein family ^28^. Thus, the ATR-FTIR signal arises from the glycoprotein that present these biomarkers, rather than from the isolated epitopes. Comparing across spectra, CA125 exhibited the strongest Amide I and Amide II contributions, followed by CA15-3, CEA, and AFP. This ordering is consistent with the relative molecular sizes of the biomarkers: CA125 (MUC16) is by far the largest molecule in the panel, with CA15-3 (MUC1) also possessing a large protein backbone. At comparable surface coverage, these larger mucins present a greater number of peptide bonds, leading to stronger amide band intensities. In contrast, the smaller, more globular glycoproteins AFP and CEA contribute fewer peptide bonds per molecule, resulting in weaker amide band signals.

**Figure 2.**
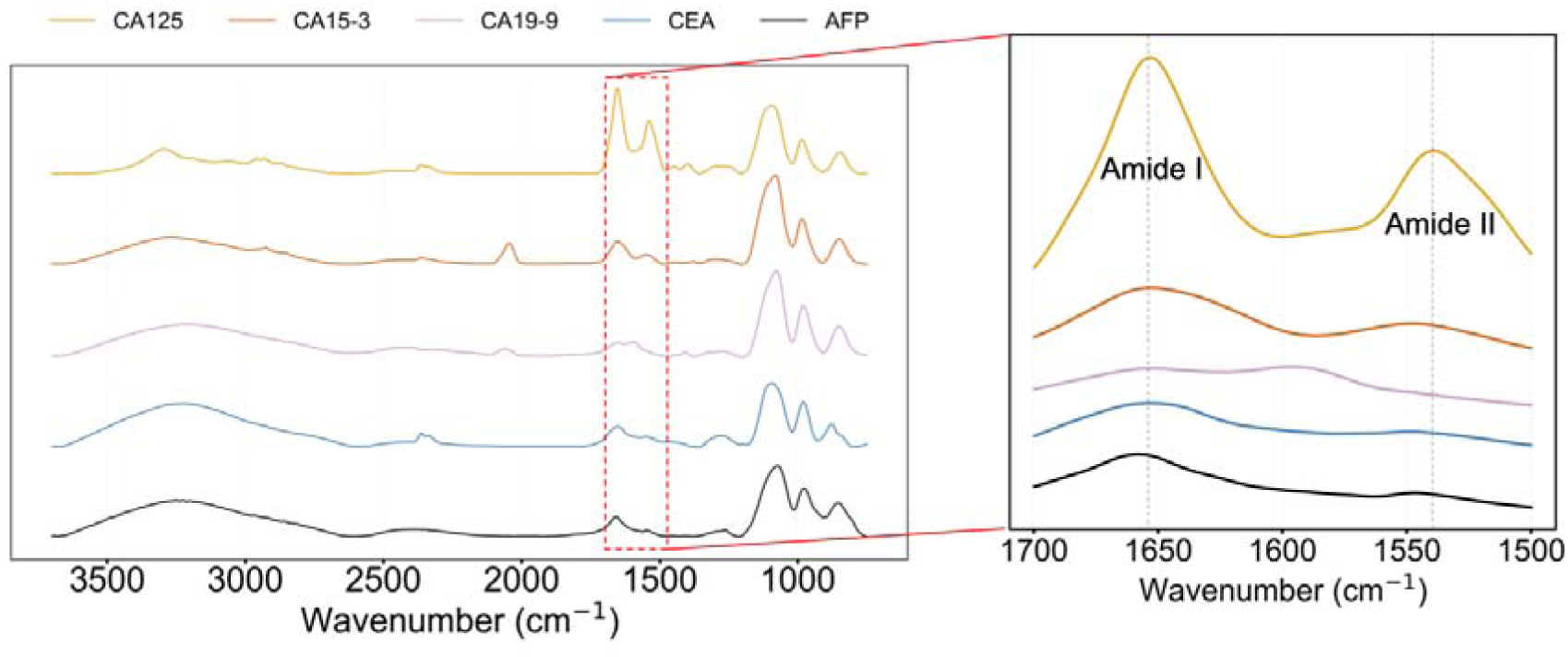
Averaged ATR-FTIR spectra of CA125, CA15-3, CA19-9, CEA and AFP. Inset shows the zoomed in region for the Amide I and Amide II regions.

Relative shifts in the Amide I and Amide II peak positions were also observed across biomarkers. Amide I positions ranged from 1653 cm^−1^ for CA125 and CA15-3, to 1655 cm^−1^ for CEA and 1658 cm^−1^ for AFP. The Amide II band showed greater variation, with CA125 appearing at 1539 cm^−1^, CA15-3 at 1545 cm^−1^, and both CEA and AFP at 1548 cm^−1^. Notably, these shifts follow a protein family trend, with the mucin glycoproteins (CA125, CA15-3) producing Amide peaks at lower wavenumbers relative to the oncofetal glycoproteins (CEA, AFP). Given that Amide I is sensitive to protein secondary structure and Amide II to backbone environment, these shifts likely reflect underlying differences in protein folding and glycosylation between the two families, although definitive assignment of secondary structure contributions would require decomposition of the Amide I band ^29^.

CA19-9 was the exception in this analysis, where the characteristic Amide I and Amide II bands observed in the other biomarkers were reduced or absent. CA19-9 is a tetrasaccharide epitope that can be carried by multiple different proteins ^30^, and its spectrum is influenced by the composition of the sugar chains, which reduces the relative contributions from protein-associated vibrational modes. A weak band near ∼1655 cm^−1^ was observed, which falls within the Amide I region, however, given the oligosaccharide composition of CA19-9, this feature may also carry contributions from C=O stretching modes of carbohydrate residues, and cannot be assigned to protein structure alone. The Amide II band, which arises more specifically from N-H bending and C-N stretching of the peptide backbone, was not as clearly discernible. Additionally, an absorption at ∼1605 cm^−1^ was present, consistent with asymmetric COO^−^ stretching ^25^, likely arising from the carboxylate group of sialic acid.

Taken together, these findings demonstrate that each biomarker produces a distinct ATR-FTIR spectral profile reflecting differences in protein composition, molecular size and glycan structure.

### PCA of cancer biomarkers

PCA was performed on the spectra of all biomarkers across three wavenumber ranges: the full spectral range (1200-3700 cm^−1^), with buffer-associated contributions excluded; the protein-specific amide region (1200-1700 cm^−1^); and the O-H/N-H stretching region (3000-3700 cm^−1^).

In the full spectral range, all five biomarkers formed clusters, with PC1 and PC2 accounting for 50.8% and 14.2% of the variance, although partial overlap was observed between CA19-9 and AFP (Figure 3a). The protein-specific amide region (1200-1700 cm^−1^) produced the clearest overall separation, with PC1 and PC2 explaining 68.0% and 12.3% of the variance (Figure 3b). The O-H/N-H stretching region (3000-3700 cm^−1^) captured a comparable proportion of variance to the amide region, with PC1 and PC2 accounting for 67.1% and 15.9%, but showed greater overlap between biomarkers, particularly between CA19-9 and AFP (Figure 3c). The clearer clustering observed in the 1200-1700 cm^−1^ amide region may indicate that spectral features outside this range introduce additional variance from non-biological sources, such as sample hydration state or atmospheric absorption, thereby reducing separation in the score plots.

**Figure 3.**
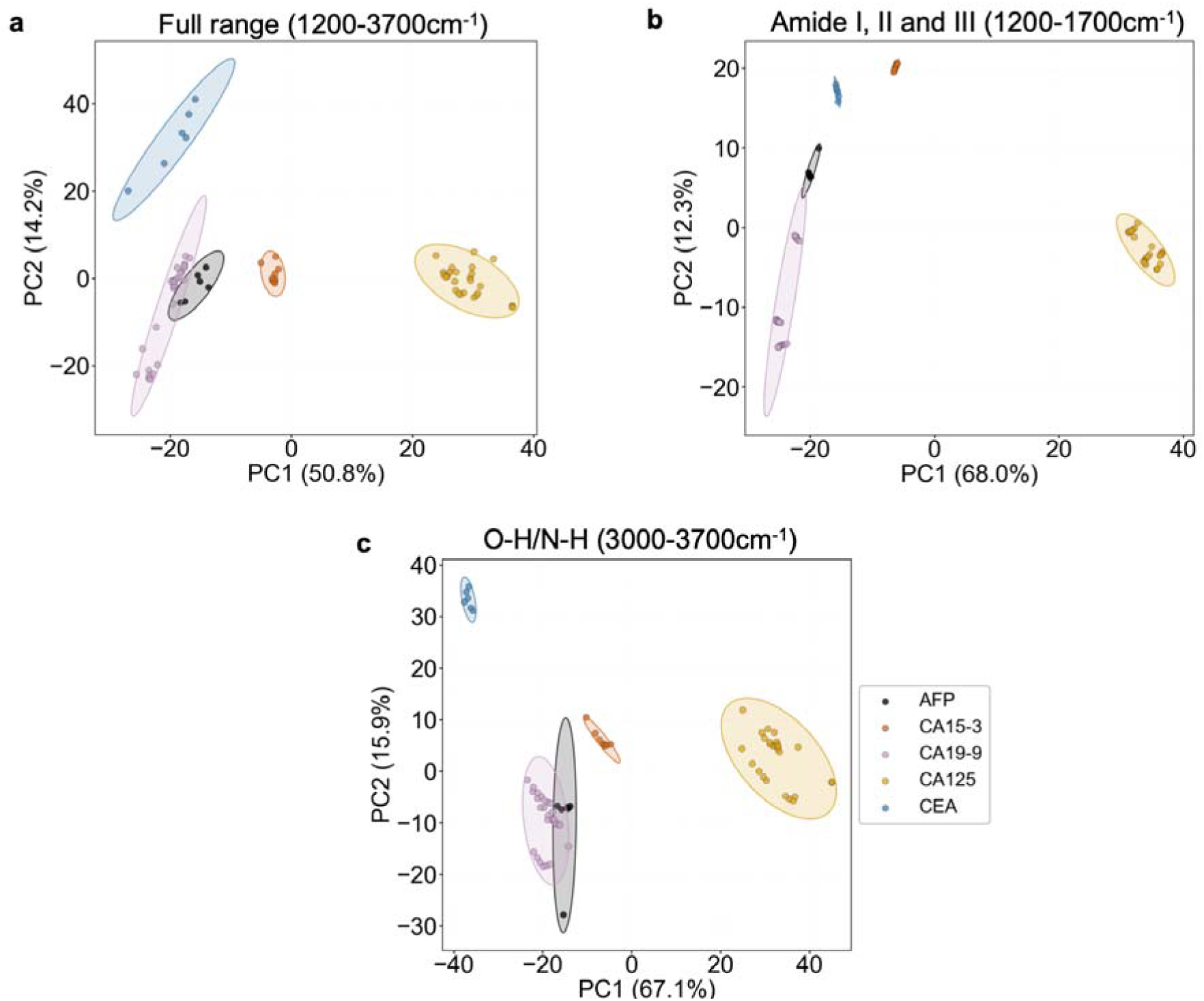
PCA of tumour biomarkers in different regions of the spectra. a) Full wavenumber range analysed (1200-3700cm^−1^) excluding the buffer contributing region. b) Amide I,II and III associated wavenumber range (1200-1700cm^−1^) and c) O-H/N-H associated wavenumber range (3000-3700cm^−1^).

Across all three ranges, CA125 showed the most pronounced separation, driven primarily along PC1, likely reflecting its amide-associated spectral contributions. CA15-3 clustered closest to CA125 and was also well separated, consistent with their shared mucin glycoprotein structure. CEA showed notable separation along PC2 in the amide region, distinguishing it from the remaining biomarkers despite its weaker amide contributions. AFP and CA19-9 clustered most closely across all ranges, which may be explained by the fact that CA19-9 is a carbohydrate epitope which has been shown to be carried by AFP, among other proteins, potentially giving rise to shared spectral features ^31^.

### Quantification of CA125 at high concentrations

We assessed concentration-dependent spectral changes in CA125 by comparing the averaged spectra of a high-concentration sample (50 kU/mL) with a lower-concentration sample (5 kU/mL). Decreasing CA125 concentration produced a reduction in the intensity of the Amide I and Amide II bands, with higher concentrations producing stronger absorbance in both regions (Figure 4). By contrast, the Amide III region and the higher-wavenumber Amide A and B bands showed less pronounced concentration-dependent variation, likely reflecting contributions from the buffer matrix to absorbance in those spectral regions. Accordingly, subsequent quantitative modelling of CA125 in PBS was restricted to the Amide I/II window (1550-1700 cm^−1^), where protein-specific features remained clearly detectable and varied systematically across the concentration range.

**Figure 4.**
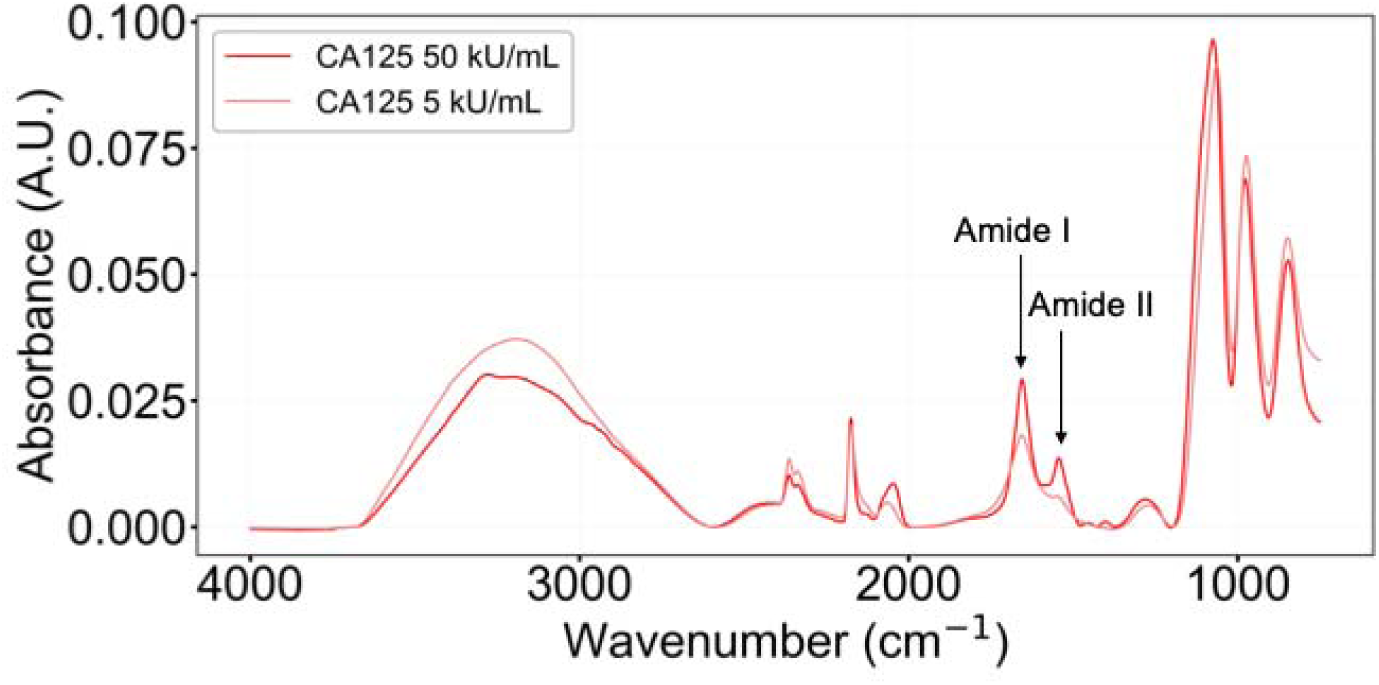
ATR-FTIR Spectra of CA125 in PBS at 50 kU/mL compared to at 5 kU/mL, showing concentration dependent changes in Amide I and Amide II band intensities.

Next, to assess whether quantitative prediction of CA125 is feasible, a PLSR model was first developed on high concentration CA125 samples. Absorbance values from the Amide I and Amide II regions across multiple CA125 concentrations were used as input variables, with the corresponding CA125 concentrations serving as the regression targets. Model optimisation was performed by selecting the number of latent variables that achieved an optimal balance between R^2^ and MSE across the cross-validation training and test sets.

Six components were identified as the optimal model complexity from a total of 25 evaluated (Figure 5a). At this point, the cross-validation MSE reaches its minimum (Figure 5b), indicating that the model captures the maximum amount of meaningful spectral variance without incorporating noise. Beyond six components, the cross-validation R^2^ begins to decline and the cross-validation MSE rises, whilst the training MSE continues to decrease. This divergence between training and cross-validation performance is characteristic of overfitting, where additional components begin to model replicate-specific spectral noise rather than the underlying concentration-dependent variation, reducing the model’s ability to generalise to unseen data.

**Figure 5.**
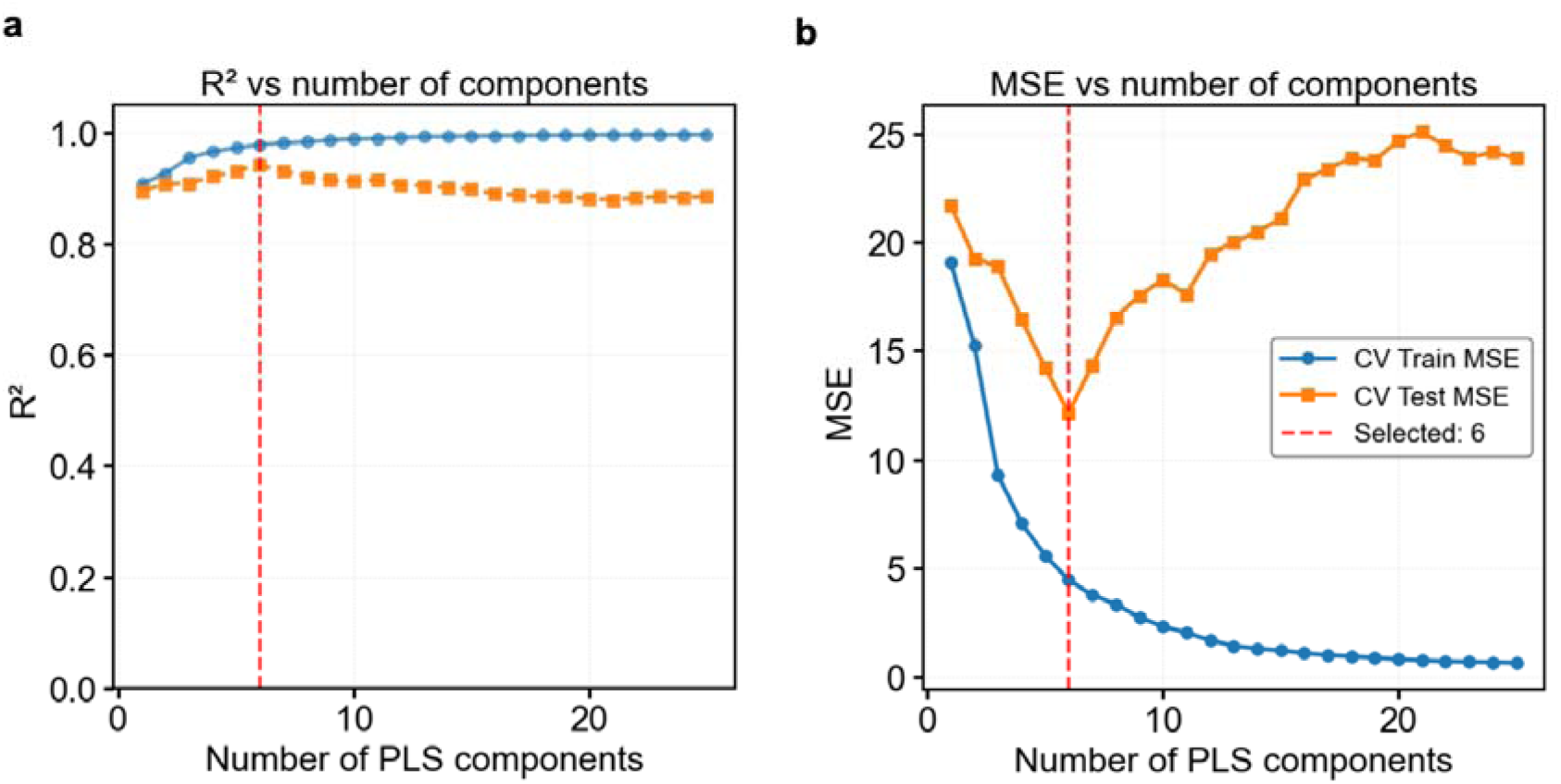
PLSR component optimisation. a) CV train (blue) and CV test (orange) R^2^ compared with number of PLS components b) CV train and CV test MSE compared with number of PLS components.

Based on this component optimisation, two PLSR models were evaluated. On the validation set, the model demonstrated strong predictive performance, with an R^2^ of 0.94 and an RMSE of 3.5 kU/mL (Figure 6a). The model generalised well to unseen data, achieving comparable performance on the independent test set with an R^2^ of 0.95 and an RMSE of 3.1 kU/mL (Figure 6b). The consistency between validation and test set metrics indicates that the model was not overfitted to the training data, and that concentration-dependent spectral variation in the Amide I/II region is sufficient to support reliable quantitative prediction of CA125 across the concentration range examined.

**Figure 6.**
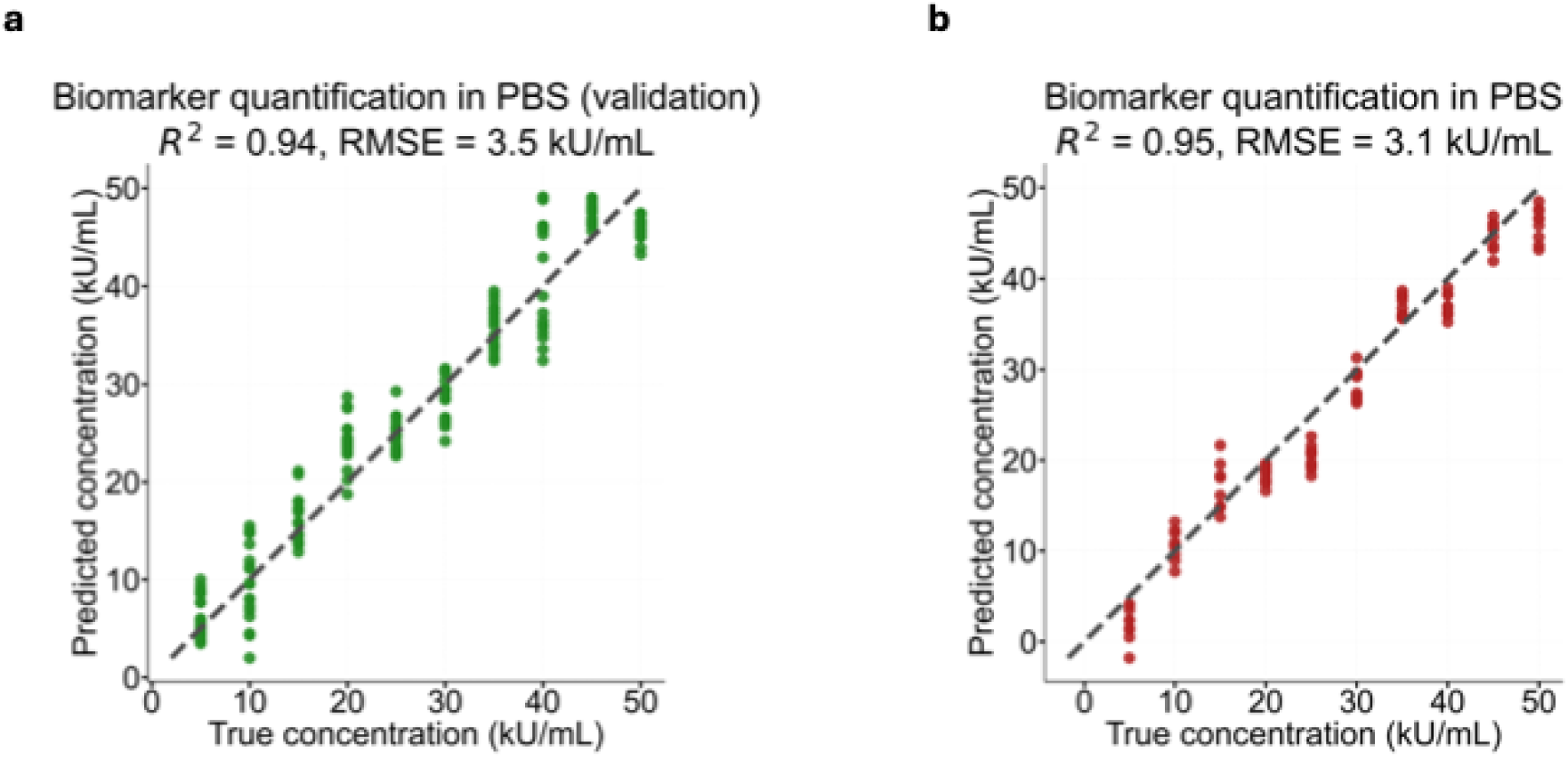
CA125 quantification at high concentrations (5 kU/mL - 50 kU/mL) using PLSR. a) PLSR model performance on validation set. b) PLSR model performance on test set.

Notably, both models demonstrated strong correlation and low prediction error across the concentration range studied. The slightly improved performance observed on the independent test set likely reflected replicate to replicate variability, as the test data comprised a completely separate experimental replicate that was not involved in model optimisation.

Overall, these results confirm that ATR-FTIR spectroscopy combined with PLSR can reliably quantify CA125 in a buffer matrix across the 5-50 kU/mL concentration range, establishing a foundation for extending the approach to more biologically complex samples.

### Analysis of human serum spiked with CA125

ATR-FTIR spectra were collected from human serum donated by a 63-year-old female and spiked with CA125 over a concentration range of 5-1000 U/mL. FTIR analysis of human serum is well established, with comprehensive peak assignments reported in the literature ^32–34^. The spectra obtained from unspiked serum (0 U/mL) and CA125-spiked serum samples (5-1000 U/mL) were dominated by the characteristic absorption profile of serum, with additional spectral changes observed following CA125 spiking (Figure 7).

**Figure 7.**
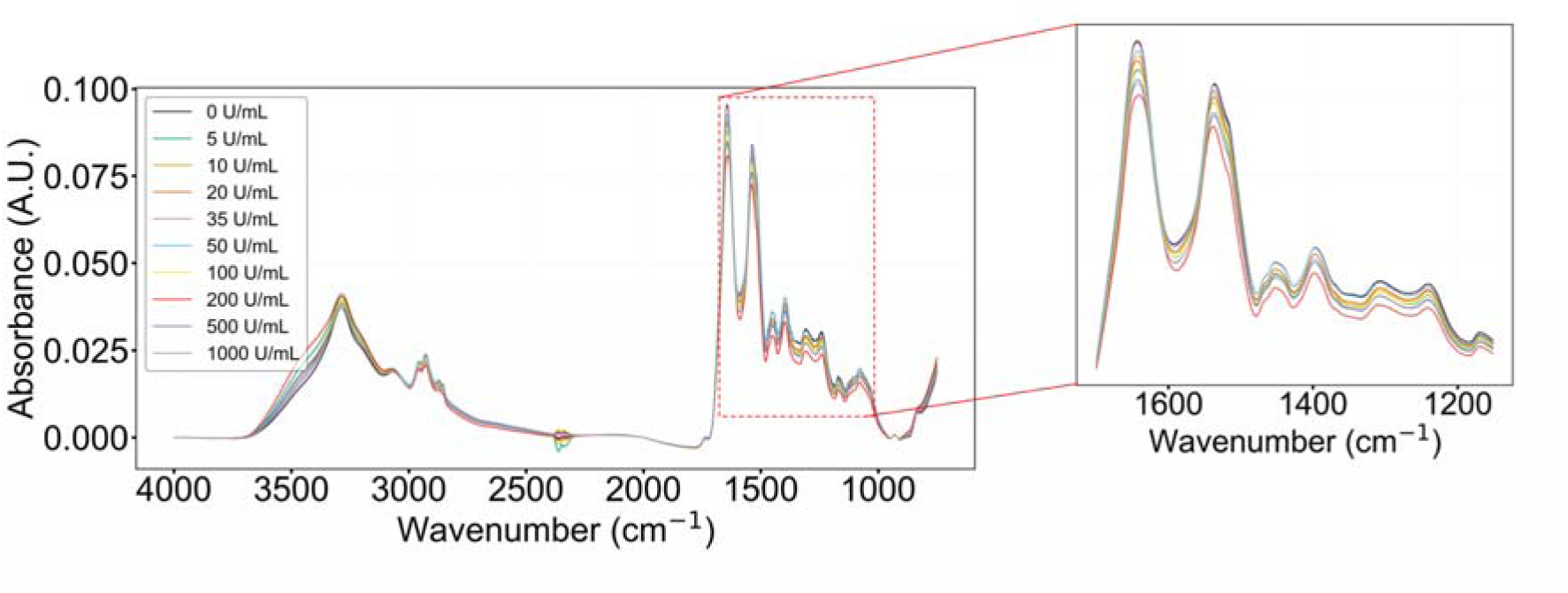
ATR-FTIR spectra of human serum spiked with CA125 at concentrations ranging from 5 to 1000 U/mL. The 0 U/mL sample corresponds to serum alone without added CA125. The zoomed inset highlights the spectral response in the 1200-1700 cm^−1^ regions.

The strong protein-associated bands were most prominent, particularly Amide I at around 1650 cm^−1^ and Amide II at around 1550 cm^−1^, consistent with the high protein content of serum. Additional absorptions were observed in the lower wavenumber region between 900 and 1200 cm^−1^, arising from C-O and C-O-C vibrations associated with carbohydrates and glycoprotein glycans, together with contributions from phosphate-containing species. Lipid-associated bands were also evident, including a carbonyl absorption near 1740 cm^−1^ and CH_2_/CH_3_ stretching bands in the 2800-3000 cm^−1^ region ^35^. In the higher wavenumber region, broad contributions between 3200 and 3600 cm^−1^ were assigned to N-H and O-H stretching vibrations, including Amide A at around 3300 cm^−1^ and hydration-related bands, while the weaker feature near 3070-3100 cm^−1^ was consistent with Amide B.

With increasing CA125 concentrations, measurable deviations in spectral intensity were observed, mostly within the protein-associated 1200-1700 cm^−1^ region. However, unlike the CA125-only spectra, in which Amide band intensities changed monotonically with concentration, the spiked serum spectra showed a non-linear response, with Amide band intensities neither consistently increasing nor decreasing as CA125 concentration increased. This might suggest that, within serum, the spectral contribution from CA125 is convoluted and influenced by the endogenous protein background, producing a more complex spectral response than that observed in purified CA125 preparations. Additionally, the non-linear spectral response may partly reflect interactions between spiked CA125 and the endogenous serum protein background. Previous studies have suggested that CA125 can exist in circulating immune complexes with immunoglobulins in serum, which may contribute to behaviour that deviates from a simple additive response ^36^.

### Quantification of CA125 in serum spiked samples

Given the concentration-dependent spectral changes observed in spiked serum, PLSR was applied to assess whether these changes could support quantitative prediction of CA125. In contrast to the PBS-based model, which focused on wavenumber regions specific to CA125 within the Amide I and Amide II bands, the serum model was applied across the full relevant spectral range (750-4000 cm^−1^) to capture all concentration associated spectral changes. To account for the wide range of CA125 concentrations (5-1000 U/mL) and to avoid excessive extrapolation leading to poor model performance, two PLSR models were constructed. One model was trained to quantify low concentrations (5-50 U/mL), while a second model targeted higher concentrations (100-1000 U/mL). In both models, the same optimisation strategy was used with the number of LVs selected to achieve an optimal balance between R^2^ and MSE in cross-validation.

Each model was assessed using its own independent test set, with performance summarised in Figure 8. For the low-concentration regime (Figure 8a), the model achieved an R^2^ of 0.77 with an RMSE of 8.4 U/mL. The estimated limit of detection (LoD) for CA125 in serum was 31 U/mL, meaning predictions below this level are close to the sensitivity limit of the method and are therefore expected to carry greater uncertainty.

**Figure 8.**
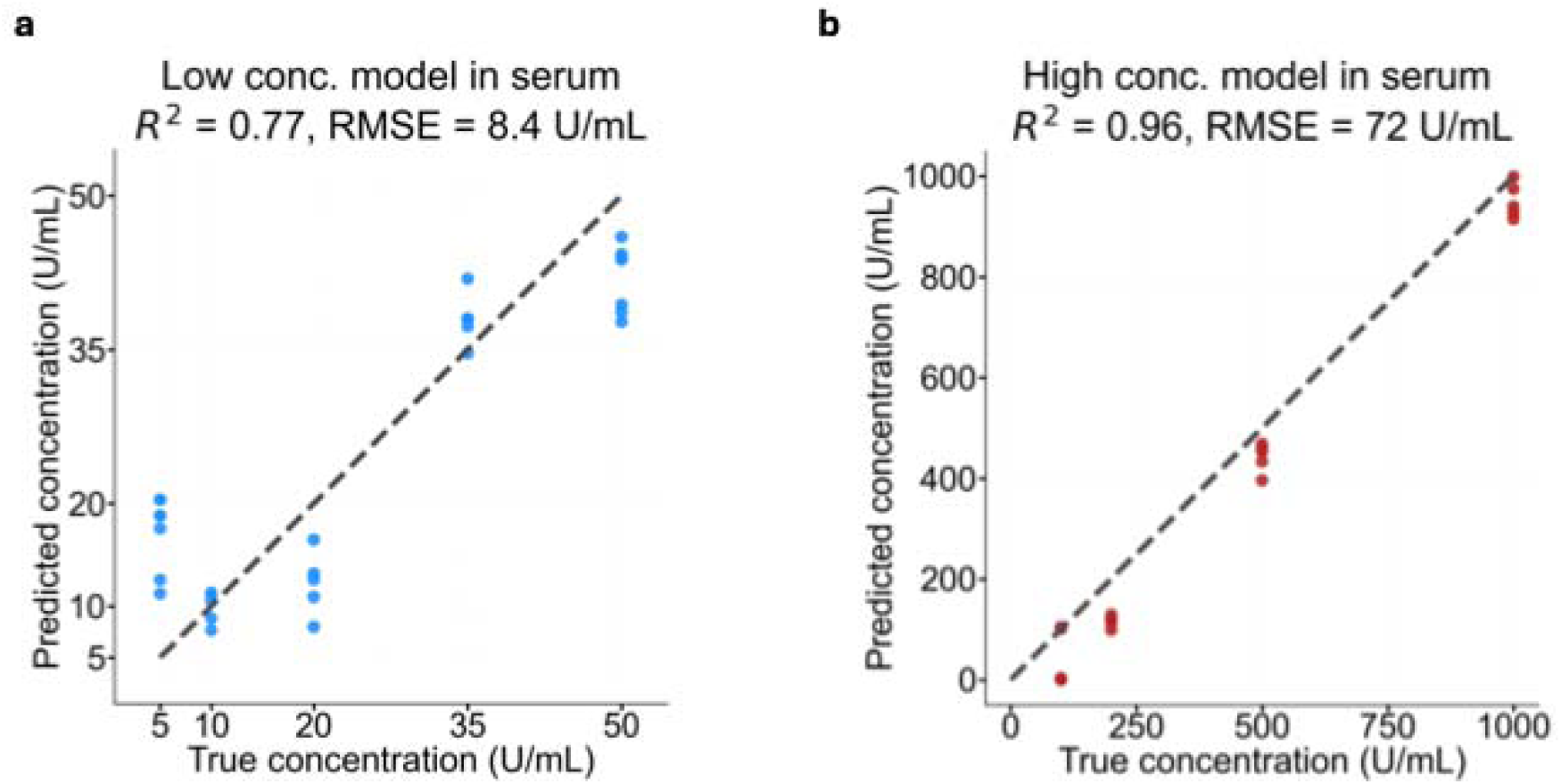
PLSR model regression CA125 in human serum across low and high concentration ranges. (a) Low concentration PLSR model trained to quantify CA125 between 5-50 U/mL. (b) High concentration PLSR model trained to quantify CA125 between 100-1000 U/mL.

For the higher concentration regime (100-1000 U/mL; Figure 8b), performance improved substantially, with an R^2^ of 0.96 and an RMSE of 72 U/mL. The largest variability was observed at the lower end of this range, particularly around 100 U/mL (with predictions spanning approximately 0-120 U/mL) and, to a lesser extent, 200 U/mL where variability persisted and underestimation was more common. In contrast, predictions for 500 U/mL and 1000 U/mL were tightly clustered near the identity line with minimal deviation, indicating that quantitative performance becomes more robust as CA125 concentration increases.

Overall, these results show that ATR-FTIR combined with PLSR enables quantification of CA125 in spiked human serum, with strongest performance at elevated concentrations and more moderate accuracy near the clinical decision threshold, while reliability improves above 35 U/mL.

### Multiclass classification of CA125 in serum spiked samples

As an alternative reformulation of biomarker analysis in human serum, a semi-quantitative multiclass classification approach was evaluated by binning serum CA125 concentrations into low (5-20 U/mL), medium (35-100 U/mL), and high (200-1000 U/mL) categories. The dataset was partitioned using a stratified, replicate-aware split, and PCA scores derived from the preprocessed training spectra were used as inputs to a regularised multinomial logistic regression classifier with balanced class weighting.

Hyperparameter optimisation across regularisation strengths and PCA component numbers (Figure 9a,b) identified C = 0.1 as the optimal regularisation, with intermediate values outperforming both strongly regularised models, which showed underfitting, and weakly regularised models, which were sensitive to model complexity. Based on validation accuracy and cross-entropy loss, 5 principal components were selected as the optimal dimensionality, corresponding to the point at which validation accuracy peaked and loss was minimised before the onset of overfitting (Figure 9c,d).

**Figure 9.**
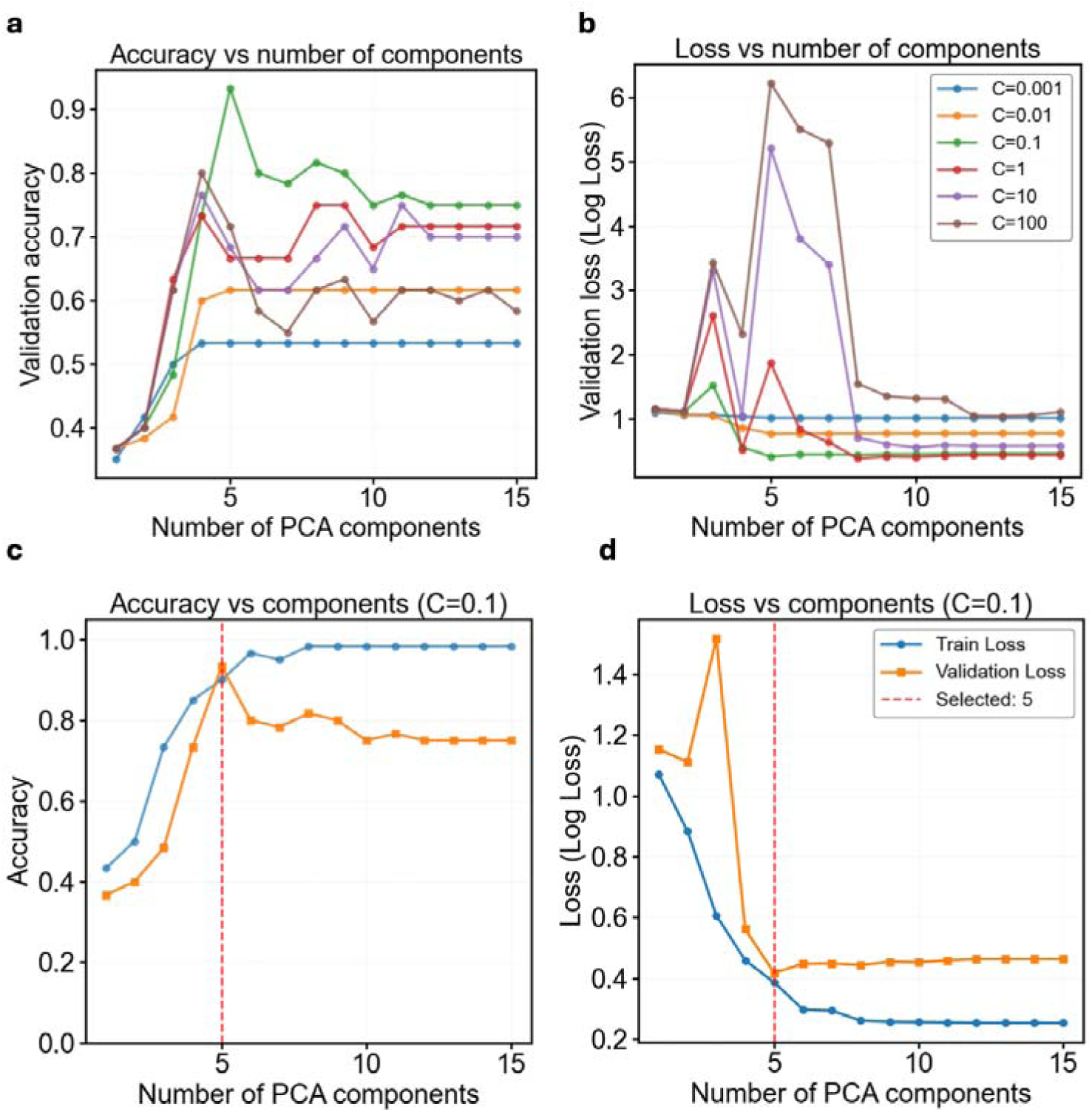
Hyperparameter optimization and feature dimensionality selection of PCA-Logistic Regression Classifier (a) Validation accuracy as a function of the number of retained PCA components (1-15) for different inverse regularization strengths (C). Models with larger C values (weaker regularization) showed greater instability across component numbers, whereas stronger regularization leads to more stable performance. The model with C = 0.1 (green curve) consistently achieves the highest validation accuracy, peaking at 5 PCA components. (b) Corresponding validation cross-entropy (log loss) curves across the same PCA component range and C values, showing a strong reduction in loss for C = 0.1 at low to moderate number of components. (c) Training (blue) and validation (orange) accuracy for the selected C = 0.1 model. The vertical dashed line marks 5 PCA components, identified as an elbow point where validation accuracy is maximized while the gap between training and validation performance remains minimal, indicating minimal overfitting. (d) Training and validation loss curves for the same model. The selection of 5 PCA components (red dashed line) representing the optimal trade-off between model complexity and generalization.

The optimised model (C = 0.1, 5 PCs) was evaluated on an independent test set, with performance summarised in Table 1 and confusion matrices shown in Figure 10. The high concentration class (200-1000 U/mL) was identified with perfect sensitivity, specificity, and precision. Misclassifications were restricted to adjacent classes, with 27% of low samples misclassified as medium and 16% of medium samples misclassified as low, likely reflecting spectral overlap near the 35 U/mL boundary. Overall, the model achieved a macro-average sensitivity of 0.86, specificity of 0.92, and precision of 0.86.

**Figure 10.**
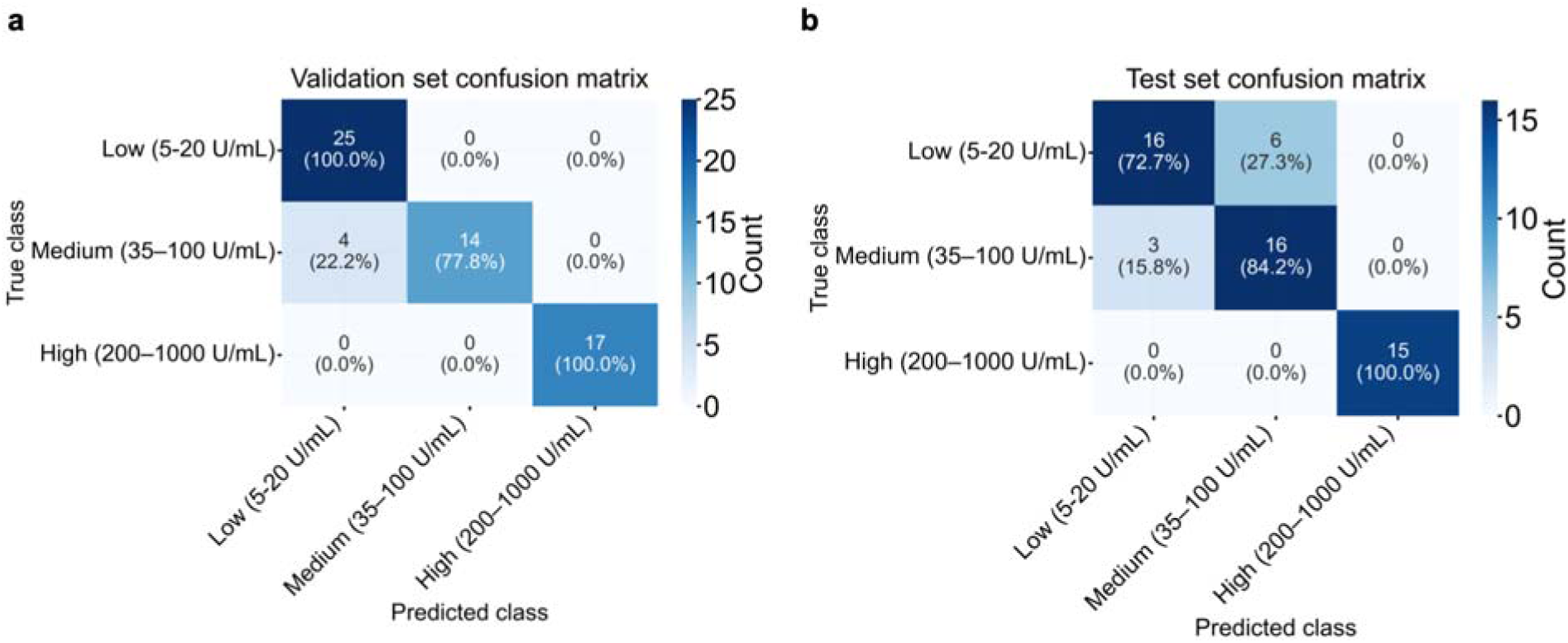
Confusion matrices showing the performance of the classification model for CA125 concentrations. Results are displayed for the (a) validation set and (b) independent test set. CA125 levels are categorized into three groups: Low (5-20 U/mL), Medium (35-100 U/mL), and High (200-1000 U/mL).

**Table 1:** PCA-LR performance metrics for serum CA125 multiclass classification.

| Class Name | Sensitivity | Specificity | Precision |
| --- | --- | --- | --- |
| Low (5-20 U/mL) | 0.73 | 0.91 | 0.84 |
| Medium (35-100 U/mL) | 0.84 | 0.84 | 0.73 |
| High (200 -1000 U/mL) | 1.00 | 1.00 | 1.00 |
| Macro average | 0.86 | 0.92 | 0.86 |

These results demonstrate that reformulating CA125 quantification as a multiclass classification problem improves clinical interpretability, particularly at concentrations near the diagnostic threshold where continuous regression showed the greatest uncertainty. The ability to reliably identify elevated CA125 levels with perfect classification performance, combined with robust discrimination above the 35 U/mL clinical cut-off, supports the potential utility of this approach as a rapid, label-free triage tool. While misclassifications between adjacent low and medium classes highlight the inherent spectral challenge near the threshold, the low false-positive rate and high specificity are encouraging from a clinical screening perspective.

## Conclusions

This study demonstrates that ATR-FTIR spectroscopy combined with machine learning has potential as a rapid, reagent-free platform for tumour biomarker analysis. Although previous ATR-FTIR studies have shown promise for cancer detection, most have framed the task as binary classification of disease presence or absence in biofluids such as whole blood or tissue. In contrast, this work addresses the less explored but clinically important problem of quantitative biomarker measurement, targeting a specific protein across a concentration range aligned with clinical decision thresholds. Moreover, rather than focusing on a single biomarker, we demonstrate spectral separability across a panel of five clinically relevant proteins using PCA within the protein-associated region (1200-1700 cm^−1^), which to our knowledge has not previously been reported using ATR-FTIR, indicating that the analytical framework is not restricted to a single target. Building on this, regression modelling of CA125, first in PBS and then in human serum, showed that quantitative prediction is achievable across a clinically relevant concentration range. Performance was strongest at elevated concentrations (R^2^ = 0.96), while a semi-quantitative classification approach improved interpretability near the diagnostic threshold. Notably, the high-concentration class associated with elevated ovarian cancer risk was identified with 100% classification accuracy, and where misclassifications occurred they were confined to lower concentration ranges where immediate clinical intervention is less likely. Taken together, these findings represent a step towards moving the field beyond binary disease classification and into a quantitative analytical framework for estimating cancer biomarker concentration in clinically relevant samples. Translation to routine clinical use will require evaluation in larger and more diverse patient cohorts, alongside further optimisation of spectral acquisition, model calibration, and analytical validation.

## Data Availability

All data supporting this study are available from the University of Southampton repository at: https://eprints.soton.ac.uk/

## Notes

The authors declare no competing financial interest.

## Acknowledgements

This work was supported by the UK Engineering and Physical Sciences Research Council (EPSRC grant EP/S03109X/1). R.F thanks EPSRC DTP PhD studentship. We thank Breast Cancer Now for funding this work as part of Programme Funding to the Breast Cancer Now Toby Robins Research Centre. S-J.S. is a Lister Institute Prize Fellow.

## References

(1) Global cancer burden growing, amidst mounting need for services. https://www.who.int/news/item/01-02-2024-global-cancer-burden-growing--amidst-mounting-need-for-services (accessed 2026-02-01).

(2) Stefan, D. C.; Tang, S. Addressing Cancer Care in Low- to Middle-Income Countries: A Call for Sustainable Innovations and Impactful Research. BMC Cancer 2023, 23 (1), 756. 10.1186/s12885-023-11272-9.

(3) Biomarkers in Cancer Detection, Diagnosis, and Prognosis. https://www.mdpi.com/1424-8220/24/1/37 (accessed 2026-02-01).

(4) Liang, S.-L.; Chan, D. W. Enzymes and Related Proteins as Cancer Biomarkers: A Proteomic Approach. Clin. Chim. Acta Int. J. Clin. Chem. 2007, 381 (1), 93–97. 10.1016/j.cca.2007.02.017.

(5) Liu, Y.-J.; Wang, C. A Review of the Regulatory Mechanisms of Extracellular Vesicles-Mediated Intercellular Communication. Cell Commun. Signal. 2023, 21 (1), 77. 10.1186/s12964-023-01103-6.

(6) Seo, J. H.; Shin, S. H.; Woo, H. R.; An, Y. R.; Youn, A. H.; Kim, S. Y.; Ki, M.-R.; Pack, S. P. Protein and Peptide in Cancer Research: From Biomarker to Biotherapeutics. Cancers 2025, 17 (18), 3031. 10.3390/cancers17183031.

(7) Scholler, N.; Urban, N. CA125 in Ovarian Cancer. Biomark. Med. 2007, 1 (4), 513–523. 10.2217/17520363.1.4.513.

(8) Duffy, M. J.; Evoy, D.; McDermott, E. W. CA 15-3: Uses and Limitation as a Biomarker for Breast Cancer. Clin. Chim. Acta 2010, 411 (23), 1869–1874. 10.1016/j.cca.2010.08.039.

(9) Luo, G.; Jin, K.; Deng, S.; Cheng, H.; Fan, Z.; Gong, Y.; Qian, Y.; Huang, Q.; Ni, Q.; Liu, C.; Yu, X. Roles of CA19-9 in Pancreatic Cancer: Biomarker, Predictor and Promoter. Biochim. Biophys. Acta BBA - Rev. Cancer 2021, 1875 (2), 188409. 10.1016/j.bbcan.2020.188409.

(10) Sauzay, C.; Petit, A.; Bourgeois, A.-M.; Barbare, J.-C.; Chauffert, B.; Galmiche, A.; Houessinon, A. Alpha-Foetoprotein (AFP): A Multi-Purpose Marker in Hepatocellular Carcinoma. Clin. Chim. Acta 2016, 463, 39–44. 10.1016/j.cca.2016.10.006.

(11) Nicholson, B. D.; Shinkins, B.; Pathiraja, I.; Roberts, N. W.; James, T. J.; Mallett, S.; Perera, R.; Primrose, J. N.; Mant, D. Blood CEA Levels for Detecting Recurrent Colorectal Cancer - Nicholson, BD - 2015 | Cochrane Library.

(12) Serum CEA Levels in 49 Different Types of Cancer and Noncancer Diseases. In Progress in Molecular Biology and Translational Science; Academic Press, 2019; Vol. 162, pp 213–227. 10.1016/bs.pmbts.2018.12.011.

(13) Mahadevarao Premnath, S.; Zubair, M. Laboratory Evaluation of Tumor Biomarkers. In StatPearls; StatPearls Publishing: Treasure Island (FL), 2025.

(14) Rusling, J. F.; Kumar, C. V.; Gutkind, J. S.; Patel, V. Measurement of Biomarker Proteins for Point-of-Care Early Detection and Monitoring of Cancer. The Analyst 2010, 135 (10), 2496–2511. 10.1039/c0an00204f.

(15) Razmi, N.; Hasanzadeh, M. Current Advancement on Diagnosis of Ovarian Cancer Using Biosensing of CA 125 Biomarker: Analytical Approaches. TrAC Trends Anal. Chem. 2018, 108, 1–12. 10.1016/j.trac.2018.08.017.

(16) Beyond ELISA: the future of biomarker validation. Drug Target Review. https://www.drugtargetreview.com/article/154316/beyond-elisa-the-future-of-biomarker-validation/ (accessed 2026-02-01).

(17) Naseer, K.; Ali, S.; Qazi, J. ATR-FTIR Spectroscopy as the Future of Diagnostics: A Systematic Review of the Approach Using Bio-Fluids. Appl. Spectrosc. Rev. 2021, 56 (2), 85–97. 10.1080/05704928.2020.1738453.

(18) Butler, H. J.; Brennan, P. M.; Cameron, J. M.; Finlayson, D.; Hegarty, M. G.; Jenkinson, M. D.; Palmer, D. S.; Smith, B. R.; Baker, M. J. Development of High-Throughput ATR-FTIR Technology for Rapid Triage of Brain Cancer. Nat. Commun. 2019, 10 (1), 4501. 10.1038/s41467-019-12527-5.

(19) Sitnikova, V. E.; Kotkova, M. A.; Nosenko, T. N.; Kotkova, T. N.; Martynova, D. M.; Uspenskaya, M. V. Breast Cancer Detection by ATR-FTIR Spectroscopy of Blood Serum and Multivariate Data-Analysis. Talanta 2020, 214, 120857. 10.1016/j.talanta.2020.120857.

(20) Giamougiannis, P.; Morais, C. L. M.; Rodriguez, B.; Wood, N. J.; Martin-Hirsch, P. L.; Martin, F. L. Detection of Ovarian Cancer (± Neo-Adjuvant Chemotherapy Effects) via ATR-FTIR Spectroscopy: Comparative Analysis of Blood and Urine Biofluids in a Large Patient Cohort. Anal. Bioanal. Chem. 2021, 413 (20), 5095–5107. 10.1007/s00216-021-03472-8.

(21) Dong, L.; Sun, X.; Chao, Z.; Zhang, S.; Zheng, J.; Gurung, R.; Du, J.; Shi, J.; Xu, Y.; Zhang, Y.; Wu, J. Evaluation of FTIR Spectroscopy as Diagnostic Tool for Colorectal Cancer Using Spectral Analysis. Spectrochim. Acta. A. Mol. Biomol. Spectrosc. 2014, 122, 288–294. 10.1016/j.saa.2013.11.031.

(22) Charkhchi, P.; Cybulski, C.; Gronwald, J.; Wong, F. O.; Narod, S. A.; Akbari, M. R. CA125 and Ovarian Cancer: A Comprehensive Review. Cancers 2020, 12 (12), 3730. 10.3390/cancers12123730.

(23) Little, T. A. Method Validation Essentials, Limit of Blank, Limit of Detection, and Limit of Quantitation | BioPharm International. https://www.biopharminternational.com/view/method-validation-essentials-limit-blank-limit-detection-and-limit-quantitation (accessed 2026-03-27).

(24) ATR-FTIR Spectroscopy and Spectroscopic Imaging of Proteins. In Vibrational Spectroscopy in Protein Research; Academic Press, 2020; pp 1–22. 10.1016/B978-0-12-818610-7.00001-3.

(25) Movasaghi, Z.; Rehman, S.; ur Rehman, Dr. I. Fourier Transform Infrared (FTIR) Spectroscopy of Biological Tissues. Appl. Spectrosc. Rev. 2008, 43 (2), 134–179. 10.1080/05704920701829043.

(26) Bouanene, H.; Miled, A. Conflicting Views on the Molecular Structure of the Cancer Antigen CA125/MUC16. Dis. Markers 2010, 28 (6), 918457. 10.3233/DMA-2010-0719.

(27) Hogendorf, P.; Skulimowski, A.; Durczyński, A.; Kumor, A.; Poznańska, G.; Oleśna, A.; Rut, J.; Strzelczyk, J. A Panel of CA19-9, Ca125, and Ca15-3 as the Enhanced Test for the Differential Diagnosis of the Pancreatic Lesion. Dis. Markers 2017, 2017, 8629712. 10.1155/2017/8629712.

(28) Zaidi, S. K.; Frietze, S. E.; Gordon, J. A.; Heath, J. L.; Messier, T.; Hong, D.; Boyd, J. R.; Kang, M.; Imbalzano, A. N.; Lian, J. B.; Stein, J. L.; Stein, G. S. Bivalent Epigenetic Control of Oncofetal Gene Expression in Cancer. Mol. Cell. Biol. 2017, 37 (23), e00352–17. 10.1128/MCB.00352-17.

(29) Barth, A. Infrared Spectroscopy of Proteins. Biochim. Biophys. Acta BBA - Bioenerg. 2007, 1767 (9), 1073–1101. 10.1016/j.bbabio.2007.06.004.

(30) Nakisa, A.; Sempere, L. F.; Chen, X.; Qu, L. T.; Woldring, D.; Crawford, H. C.; Huang, X. TumorCAssociated Carbohydrate Antigen 19–9 (CA 19–9), a Promising Target for AntibodyCBased Detection, Diagnosis, and Immunotherapy of Cancer. Chemmedchem 2024, 19 (24), e202400491. 10.1002/cmdc.202400491.

(31) Nagata, A.; Komoda, T. [CA 19-9 like epitope(s) on carcinoembryonic antigen molecules]. Radioisotopes 1989, 38 (12), 513–515. 10.3769/radioisotopes.38.12_513.

(32) Bel’skaya, L. V.; Sarf, E. A.; Solomatin, D. V. Application of FTIR Spectroscopy for Quantitative Analysis of Blood Serum: A Preliminary Study. Diagnostics 2021, 11 (12), 2391. 10.3390/diagnostics11122391.

(33) Spalding, K.; Bonnier, F.; Bruno, C.; Blasco, H.; Board, R.; Benz-de Bretagne, I.; Byrne, H. J.; Butler, H. J.; Chourpa, I.; Radhakrishnan, P.; Baker, M. J. Enabling Quantification of Protein Concentration in Human Serum Biopsies Using Attenuated Total Reflectance – Fourier Transform Infrared (ATR-FTIR) Spectroscopy. Vib. Spectrosc. 2018, 99, 50–58. 10.1016/j.vibspec.2018.08.019.

(34) Li, Y.; Li, F.; Yang, X.; Guo, L.; Huang, F.; Chen, Z.; Chen, X.; Zheng, S. Quantitative Analysis of Glycated Albumin in Serum Based on ATR-FTIR Spectrum Combined with SiPLS and SVM. Spectrochim. Acta. A. Mol. Biomol. Spectrosc. 2018, 201, 249–257. 10.1016/j.saa.2018.05.022.

(35) Frost, O. Analyzing Biofluids with ATR-FTIR Spectroscopy. AZoM. https://www.azom.com/article.aspx?ArticleID=22517 (accessed 2026-02-02).

(36) Cramer, D. W.; O’Rourke, D. J.; Vitonis, A. F.; Matulonis, U. A.; DiJohnson, D. A.; Sluss, P. M.; Crum, C. P.; Liu, B. C.-S. CA125 Immune Complexes in Ovarian Cancer Patients with Low CA125 Concentrations. Clin. Chem. 2010, 56 (12), 1889–1892. 10.1373/clinchem.2010.153122.

